# Structure of the N-terminal didomain d1_d2 of the Thrombospondin type-1 domain-containing 7A

**DOI:** 10.1101/2023.05.03.539264

**Authors:** Alice Bochel, Simon A. Mortensen, Larissa Seifert, Felicitas E. Hengel, Cy M. Jeffries, Grzegorz Chojnowski, Oliver Kretz, Tobias B. Huber, Nicola M. Tomas, Matthias Wilmanns

**Author notes:** Correspondence: Matthias Wilmanns.

## Abstract

Thrombospondin type-1 domain-containing 7A (THSD7A) is a large extracellular protein that is found in podocyte foot processes of the kidney glomerulus. It has been established as a causative autoantigen in membranous nephropathy. Amongst the predicted 21 thrombospondin repeat domains of its extracellular segment, the highest frequency of autoimmune response has been associated with the two N-terminal domains. Here, we show that antibodies against this THSD7A segment in mice induce typical clinical and morphological signs of membranous nephropathy. The high-resolution structure of these two domains reveals a non-canonical thrombospondin repeat fold that is distinct from the established type 1 thrombospondin repeat. As it shares a conserved disulfide pattern with the canonical fold, we refer to these domains d1 and d2 as type 1A thrombospondin repeats. Both domains comprise a seven layered CC-W-PP-R-W-QQ-CC pattern, which is only partly shared by other THSD7A thrombospondin repeat domains. The two domains form a well-defined V-shaped tandem arrangement. Our findings provide crucial insight into specific structural features of these two domains that are distinct from other regions of THSD7A and hence could cause the high level of antigenicity found for these two domains.

## Introduction

Membranous nephropathy (MN) is an autoimmune glomerular disease and the leading cause of nephrotic syndrome in adults. Patients are affected by severe proteinuria due to antibody-mediated damage to the glomerular filtration barrier, which is reflected by histological changes such as thickening of the glomerular basement membrane, subepithelial deposition of immune complexes and podocyte effacement (Polanco et al., 2012, Schieppati et al., 1993, Tomas et al., 2021). The M-type phospholipase A2 receptor 1 was first identified as dominant autoantigen of MN (Beck et al., 2009). Among other potential antigens, thrombospondin type-1 domain-containing 7A (THSD7A), was detected subsequently, accounting for up to 5 % of all MN cases (Tomas et al., 2014).

THSD7A is a glycosylated, type I transmembrane protein with a large, extracellular, N-terminal region (Y59-T1606, 1547 residues excluding signal sequence), a single transmembrane helix (W1607-C1630, 23 residues) and a short cytoplasmic C-terminal region (K1631-M1657, 26 residues). In endothelial cells, THSD7A regulates cell migration and tube formation in angiogenesis (Wang et al., 2010, Kuo et al., 2011). In podocytes of the kidney glomerulus, THSD7A localizes to the slit diaphragm, the endocytic compartments and the basal membrane of foot processes, where it plays a role in podocyte adhesion to the glomerular basement membrane (Herwig et al., 2019, Gödel et al., 2015). However, its physiological function and dysfunction upon antibody exposure in MN is not yet fully understood. Recruitment of IgG antibodies to the glomerulus, antibody deposition with subsequent complement activation and membrane attack complex formation explains the podocyte damage only partially. Depleting the central complement component C3 significantly attenuates MN but does not prevent or fully rescue the histological and clinical disease phenotype (Seifert et al., 2023). This finding provokes the question for a potential antigen-related mechanism of disease progression whereby autoantibody binding could damage podocytes by disrupting THSD7A function through masking binding sites or inducing structural changes (Seifert et al., 2018). To unravel molecular disease mechanisms of THSD7A-associated MN an in-depth understanding of THSD7A function and overall structure is crucial.

The extracellular region of THSD7A has been predicted to be composed of an alternating pattern of 21 type-1 and type-2 thrombospondin repeats (TSRs) (Stoddard et al., 2019, Seifert et al., 2018). TSR domains are found in extracellular regions of transmembrane or secreted proteins, where they interact with extracellular matrix components (Resovi et al., 2014). TSRs are generally composed of small three-stranded β-sheet structures, in which the first strand tends to be irregular (Tucker, 2004). In addition, they are connected by three structurally conserved disulfide bridges, which are at different positions in type-1 and type-2 TSRs (Tan et al., 2002). TSRs have a conserved WxxW motif, which in some cases is extended to a double WxxWxxW motif. There are several reports demonstrating that this motif provides a site for tryptophan-mediated C-mannosylation (Olsen and Kragelund, 2014, Crine and Acharya, 2022). Interestingly, in most of the predicted THSD7A type-1 TSR domains, the canonical WxxW TSR motif is replaced by an elongated WxxxxW motif but any specific functional role remained enigmatic (Stoddard et al., 2019). In a recent study, we identified specific autoimmunity regions in THSD7A-ssociated MN and found the highest frequency in the two N-terminal THSD7A TSR domains d1 and d2 (Seifert et al., 2018). Here, we corroborated these findings by *in vivo* experiments with an antibody repertoire limited to d1 and d2. We further characterized the crystal structure of this highly antigenic region and unexpectedly found a V-shaped arrangement as well as a distinct overall fold due to conserved proline residues. These features may crucially contribute to the unique properties and antigenicity of the two N-terminal THSD7A domains d1 and d2 in MN.

## Results

### Antibodies against the THSD7A N-terminal extracellular domains d1_d2 induce MN in mice

To assess the biological relevance of THSD7A domains d1 and d2 as immunogenic hotspot in MN, we asked whether antibodies against d1_d2 are sufficient to cause MN in mice. To this end, we immunized mice lacking THSD7A (*Thsd7a*^*−/−*^ mice) with either the complete extracellular region or d1_d2 didomain (**Figure 1A**). THSD7A mIgG recognized multiple domains of THSD7A. Anti-d1_d2 mIgG reacted with d1_d2 and d2_d3 didomains, indicating that anti-d1_d2 antibodies do not cross-react with other thrombospondin domains in THSD7A (**Figure 1B**). Neither control mIgG nor mIgG-depleted anti-THSD7A serum reacted with THSD7A domains (**Figure 1B**).

**Figure 1:**
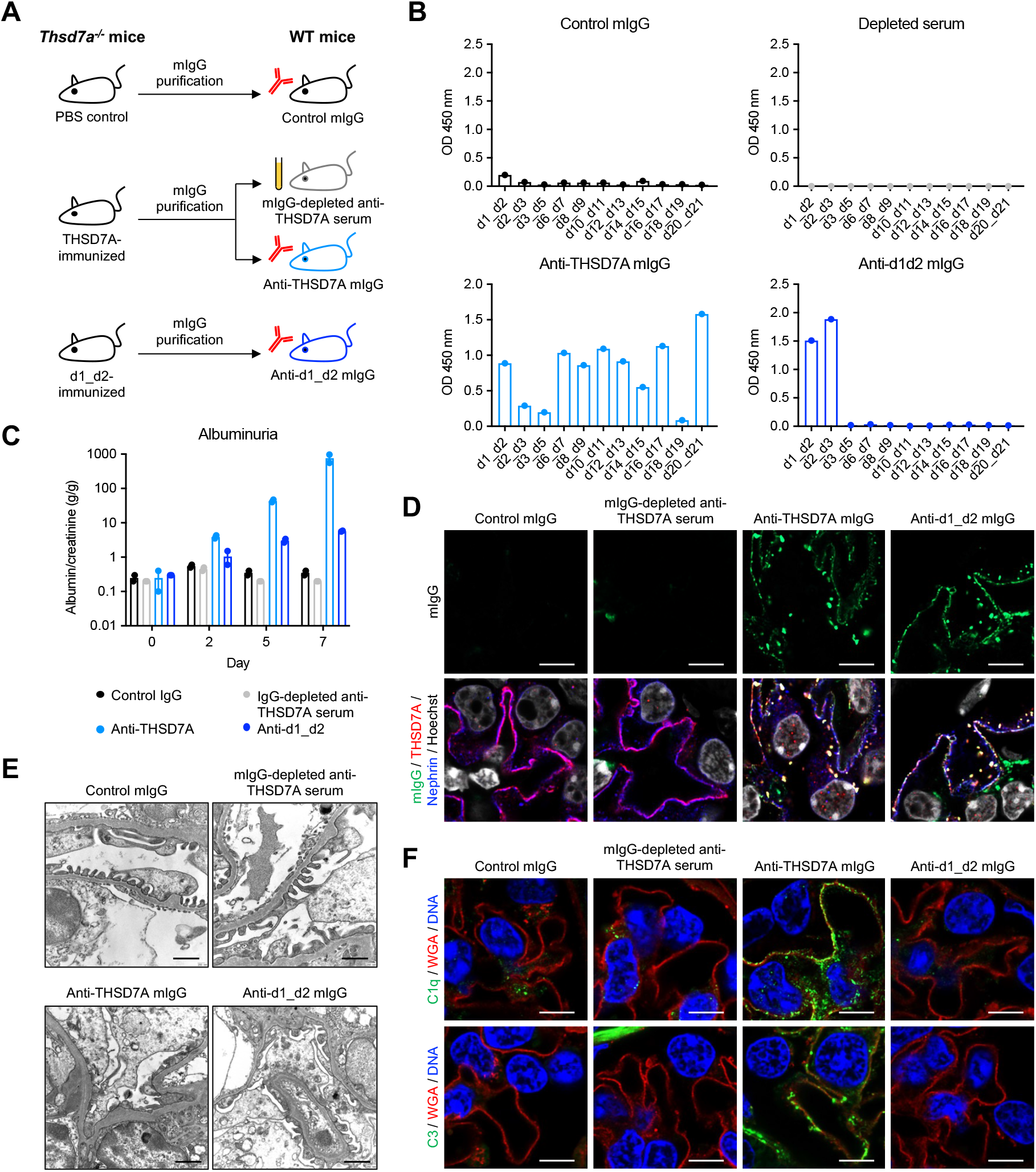
Antibodies against the d1_d2 didomain of THSD7A are sufficient to induce membranous nephropathy. (A) Experimental setup: *Thsd7a*^*−/ −*^ mice were immunized with PBS (control), full-length THSD7A, and the d1_d2 didomain of THSD7A. Murine IgG (mIgG) was then purified from the sera of immunized mice and transferred to wild-type (WT) mice. The mIgG-depleted serum from the THSD7A-immunized mice served as an additional control. (B) Domain reactivity of purified mIgG fractions and the mIgG-depleted anti-THSD7A serum. (C) Albuminuria as measured by the urinary albumin-to-creatinine ratio. (D) Immunofluorescence staining for mIgG in co-localization with THSD7A and nephrin in WT mice seven days after mIgG/serum transfer. Scale bars indicate 4 μm. (E) Electron microscopic analysis in WT mice seven days after mIgG/serum transfer. Scale bars indicate 1 μm. (F) Immunofluorescence staining for complement C1q and C3 in co-localization with wheat germ agglutinin (WGA) in WT mice seven days after mIgG/serum transfer. Scale bars indicate 4 μm.

Next, we injected WT mice each with control mIgG, mIgG-depleted anti-THSD7A serum, anti-THSD7A mIgG, and anti-d1_d2 mIgG (**Figure 1A**). Animals challenged with anti-THSD7A mIgG developed severe albuminuria (**Figure 1C**). Animals exposed to anti-d1_d2 mIgG also developed significant disease, yet to a lesser extent than anti-THSD7A mIgG-injected animals. Mice receiving anti-THSD7A mIgG or anti-d1_d2 mIgG showed typical histological signs of MN such as strong subepithelial granular deposition of mIgG (**Figure 1D**), as well as prominent podocyte foot process effacement and electron-dense deposits in electron microscopy analyses (**Figure 1E**). Interestingly, mice receiving anti-THSD7A mIgG, but not mice injected with anti-d1_d2 mIgG, also showed strong positivity for complement components C1q and C3 (**Figure 1F**), suggesting that a more restricted antibody binding to the target antigen induces less complement activation. In summary, these experiments show that antibodies against the immunodominant d1_d2 region of THSD7A induce the typical clinical and morphological signs of MN without significant activation of the complement system, pointing towards an even more important alternative molecular injury mechanism.

### Structures of THSD7A domains d1 and d2 define a distinct TSR type

The X-ray structure of human THSD7A d1_d2 di-domain was determined to 2.3 Å resolution **(Figure 2A, Supplementary Table S1)** by molecular replacement using structures of d1 and d2 predicted by AlphaFold2 (AF2) as search models. No interpretable electron density was observed for residues A48–A53 at the N-terminus of d1, as well as residues S137–E145 in the AB loop and Q192 at the C-terminus of d2. A comparison of the experimental THSD7A structure with the deposited AF2 model (UNIRPOT Q9UPZ6) reveals a reasonably correct prediction at the level of single domains d1 and d2 but the V-shaped arrangement of the two domains to each other is markedly different **(Figure 2, Supplementary Figure S1)**. Hence, reliable superimposition of the complete experimental structure of the THSD7A d1_d2 didomain with the corresponding AF2 model is not possible.

**Figure 2:**
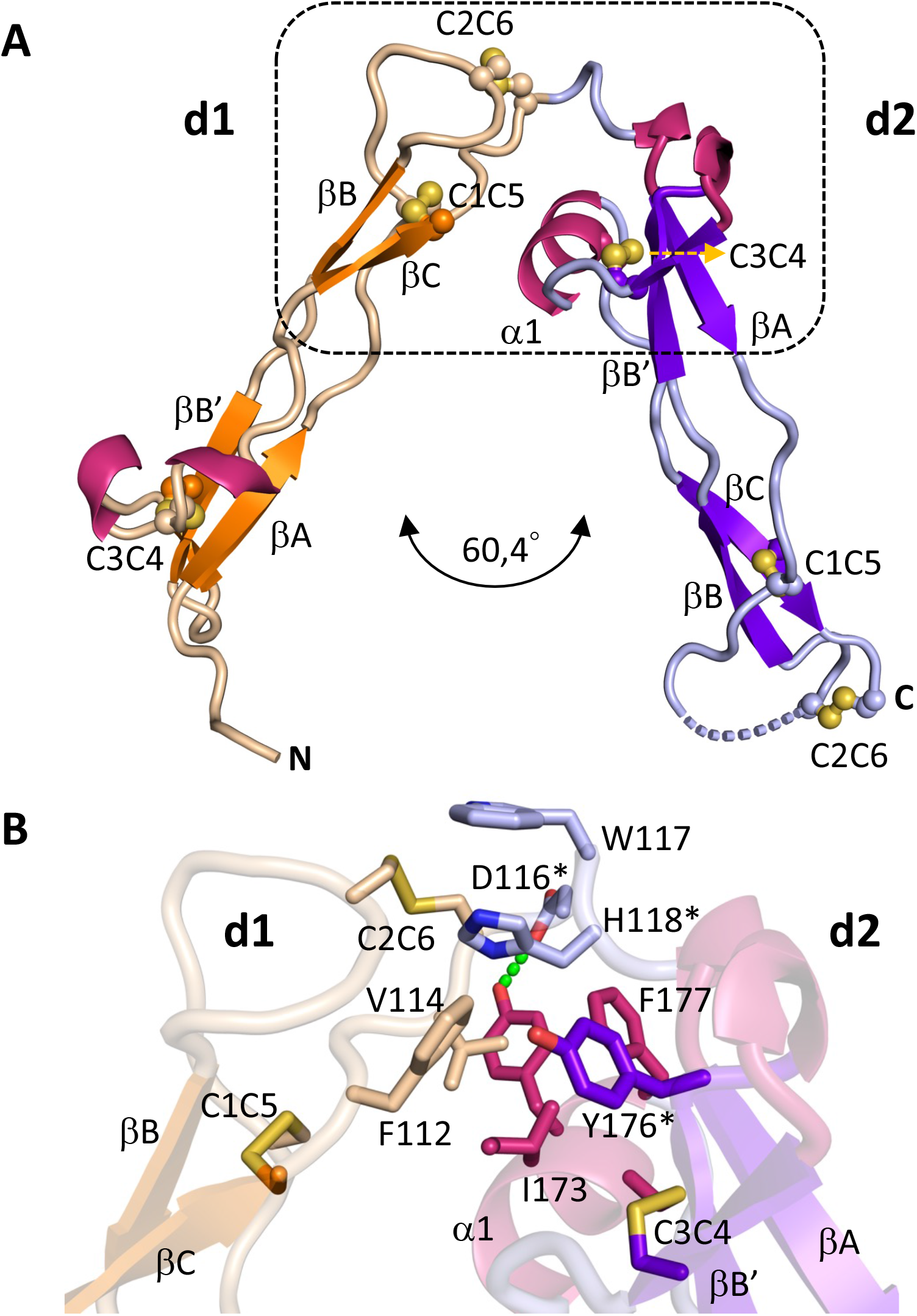
Structure of THSD7A d1_d2 didomain. Color codes: d1: β-strands, orange; connecting loops, wheat. d2: b-strands, violet, connecting loops, light blue. Helices of d1 and d2, pink. Disulfide bridges and secondary structural elements are labeled. Missing loop segment in d2 is indicated by dashed lines. A, overall V-shaped arrangement of domains d1 and d2. B, zoom into the d1_d2 interface. Interacting residues are labeled. Selected sidechain-mediated hydrogen bonds are indicated. Residues that were used to generate the d1_d2 (AAA) mutant are indicated by asterisks.

The structures of the two THSD7A domains d1 and d2 both reveal a distorted type1-1 TSR fold **(Figure 2A)**. The coordinates of both domains can be superimposed with an r.m.s.d. value of 1.33 Å for 51 matching residues out of 62 for the d1 domain and 67 residues of the d2 domain **(Figure 3A, Figure 4)**. There are three matching sequence segments, covering the three extended strand-like structures A, B, C of the type1-1 TSR fold. However, these segments only partially adopt a regular β-strand conformation **(Figure 3A-B)**. The sequence identity of the structure-based alignment is 31% **(Figure 4)**. The overall dimension of the d1 and d2 structures along the 3-stranded ABC structure from the N-terminal to the C-terminal tip is about 55 Å in length. Neither the structure of d1 nor d2, however, is superimposable with any known type-1 TSR repeat structure, despite a conserved disulfide pattern. Using the central conserved C1C5 disulfide bridge as basis for superimposition reveals that the overall bent in d1 and d2 is distinct from those observed in canonical TSR domains **(Figure 5)**. Acknowledging the conserved disulfide pattern shared with type-1 TSR domains, we have termed the fold of d1 and d2 as type 1A TSR.

**Figure 3:**
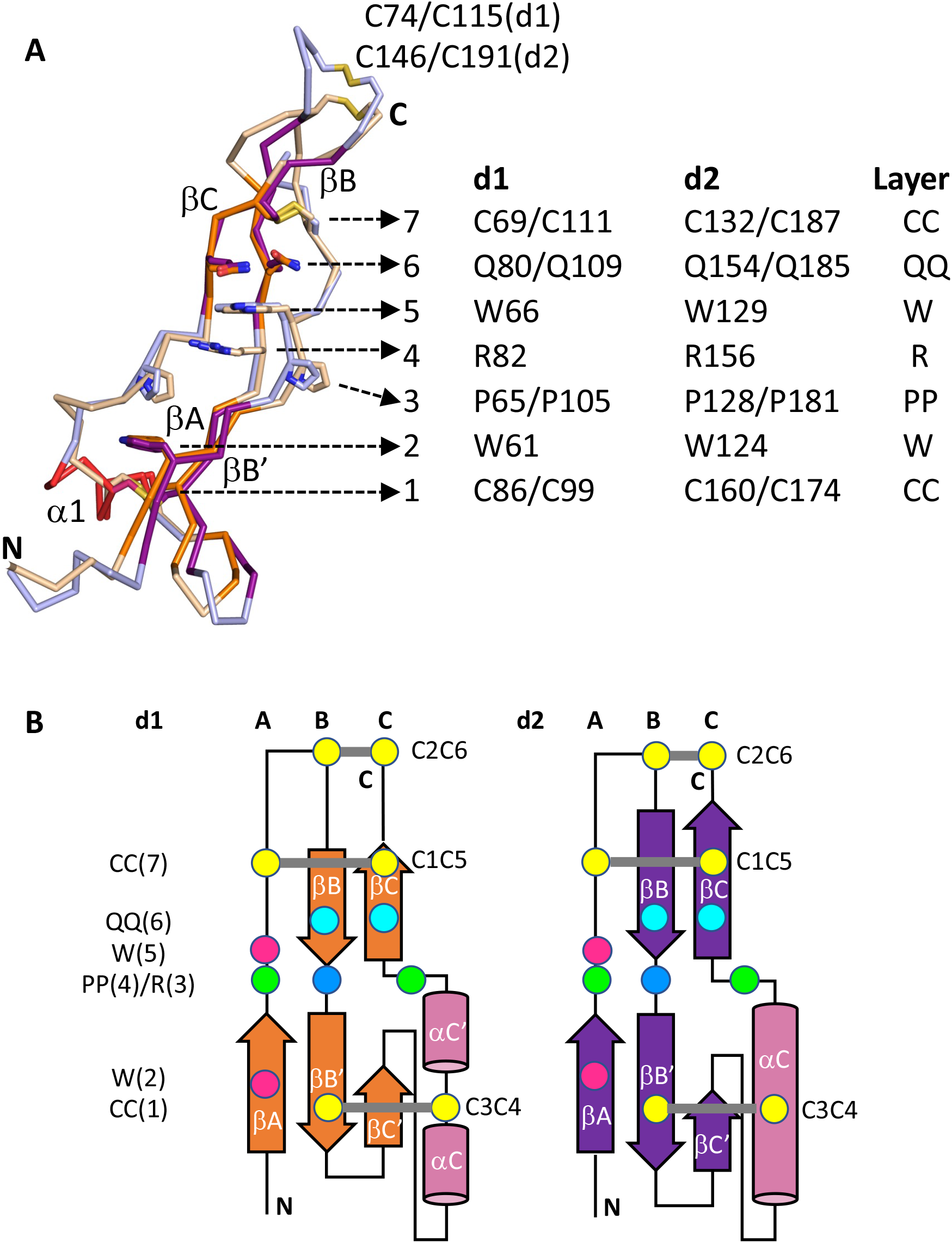
Conserved non-canonical type 1A TSR topology of THSD7A d1 and d2. A, superposition of d1 and d2. Conserved residues establishing the CC-W-PP-R-W-QQ-CC layer structure in d1 and d2 are indicated. B, topology diagrams of d1 and d2. Residues establishing the conserved CC-W-PP-R-W-QQ-CC layer structure are indicated by spheres: cysteine, yellow; tryptophan, magenta; proline, green; arginine, blue; glutamine; cyan. Colors of secondary structural elements are as in **Figure 2**.

**Figure 4:**
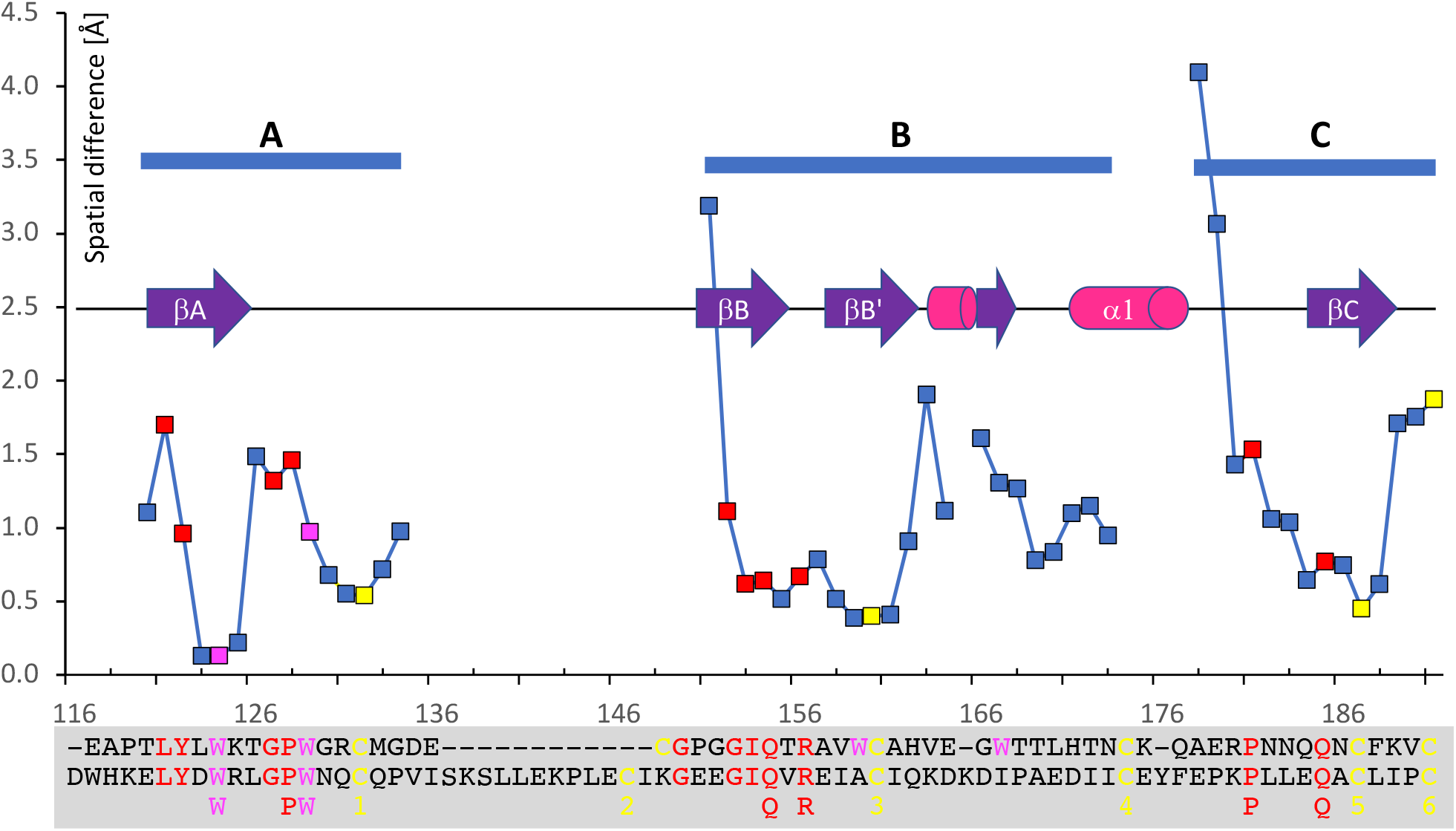
Spatial differences for each matching residue positions of superimposed d1 and d2 structural segments A, B and C. Color codes of matching residue positions: cysteine, yellow; tryptophan, magenta; other conserved residues, red. Residue numbers refer to d2 sequence. Structure-based sequence alignment is shown below. Conserved residues that contribute to CC-W-PP-R-W-QQ-CC layer structure are highlighted.

**Figure 5:**
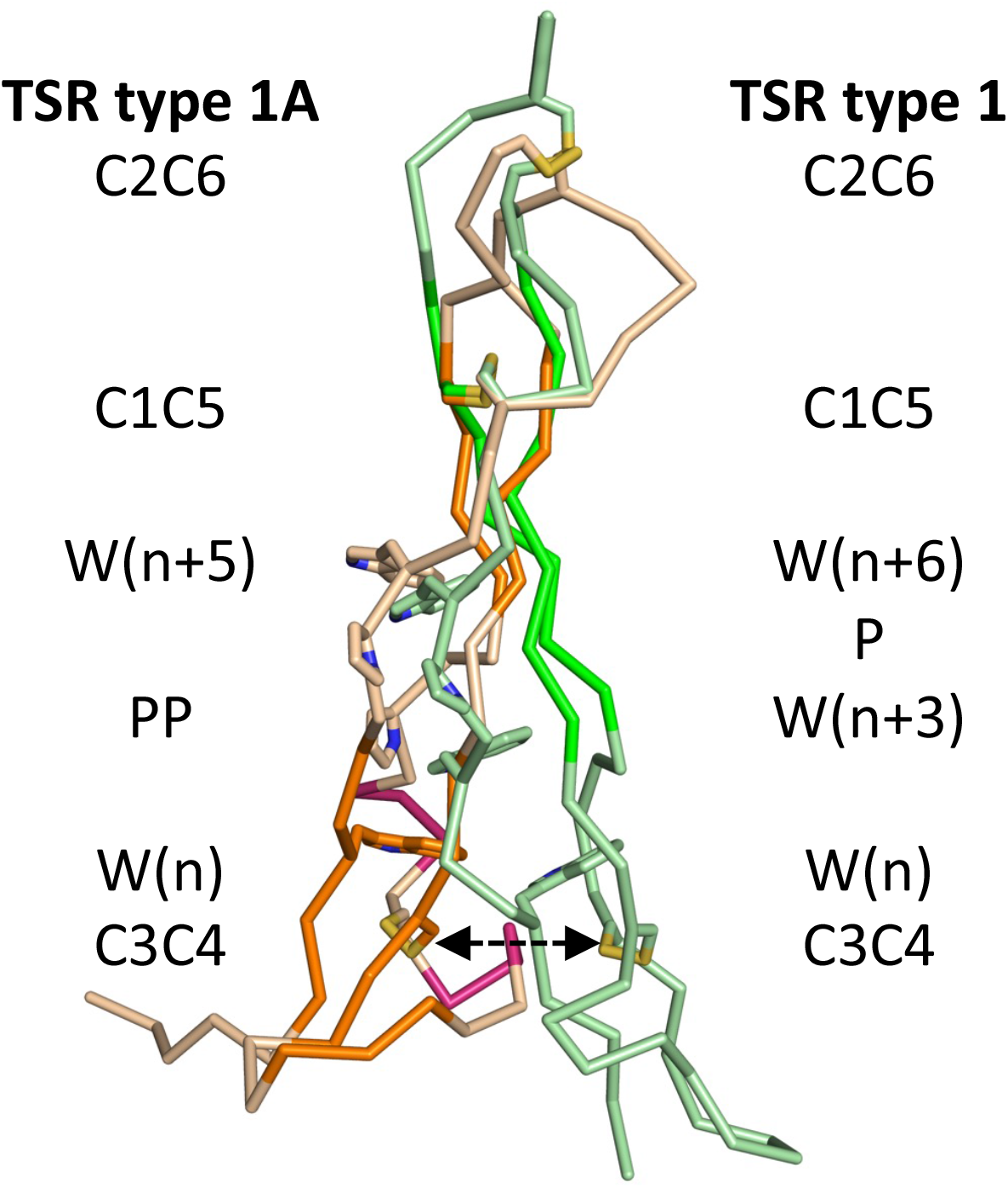
Structural superposition of the THSD7A d1 domain onto a type 1 TSR domain (PDB code: 1lsl). While an overall superposition does not provide any reliable structure-based alignment, an alignment based on the conserved C1C5 disulfide bridge, indicates deviating domain bending observed for THSD7A d1 (shown) and d2. Color codes: THSD7A as in **Figure 2**. 1lsl: b-strands, green; remaining sequence segments, pale green. Because of deviating domain bending the distance of the C3C4 disulfide bridge is about 8 Å, as indicated by a double arrow.

The sequences of d1 and d2 both have two conserved proline residues in central positions of the strand-like structures A and C, which terminates the central part of a more extended β-sheet arrangement in other type-1 TSR structures, and generates a central segment without a regular β-sheet structure in both THSD7A domains d1 and d2 **(Figure 2A, Figure 3)**. Whereas the first proline in the strand-like structure A (P65 in d1, P128 in d2) induces a local β-bulge arrangement, the second proline on the strand-like structure C (P105 in d1, P181 in d2) is oriented towards the other two strand-like structures A and B. Only in the second strand-like structure B, a proper β-strand conformation is formed before and after the central break of a regular β-sheet hydrogen bond pattern. The first one contributes to a three-stranded β-sheet with strand-like structure A and a short loop segment preceding the strand-like structure C (β-sheet AB’C’) and the second participates in a two-stranded β-sheet with strand-like structure C (β-sheet BC) **(Figure 3B)**. Strand βC is preceded by a distinct helix α1 in an orthogonal orientation to the β-sheet like structure of domains d1 and d2. While in d2 this helix has a length of 7 residues (171–177), in d1 we observed two short 310 helices instead (96–98, 101–103). There are also additional short 310 helical segments in the corresponding N-terminal tip segments of both domains d1 and d2 **(Figure 2A)**.

Despite the high level of sequence diversity between d1 and d2 mostly in the connecting loop segments at the two tips of both type-1A TSR folds, we observed a remarkable level of conservation of other residues in central positions, in addition to the conserved prolines and disulfide bridges. The WxxxxW motif is conserved as W(K/R)(L/T)GPW motif in d1 and d2 and the corresponding residues are in well matching structural positions **(Figures 3 and 4)**. Similar to findings in other type-1 TSR domain structures, the respective tryptophan side chains halfway insert into the overall β-sheet like structures **(Figures 3 and 4)**, keeping potential C-mannose attachment sites in exposed positions. In addition, we found a structurally highly conserved arginine between strands βB and βB’ (R82 in d1, R156 in d2) as well as a pair of conserved glutamines in sheet βBβC (Q80 and Q109 in d1, Q154 and Q185 in d2). Together these residues form a well-defined seven layered structure CC(1)-W(2)-PP(3)-R(4)-W(5)-QQ)6)-CC(7), where the boundaries are defined by two of three conserved disulfide bridges, C1C5 and C3C4 **(Figures 2A and 3)**. This layer structure is distinct from those observed in other type-1 TSR domain structures with characteristic tryptophan ladders (Olsen and Kragelund, 2014). In the remaining text we use the term “layer” as opposed to “ladder”, as there are contributions to it from all three extended strand segments A, B and C **(Figure 3B)**.

### V-shaped arrangement of THSD7A d1 and d2 structures

The structure of the THSD7A di-domain reveals a well-defined V-shaped C-terminal tip (d1) / N-terminal tip (d2) arrangement of the two domains, related by an angle of 60.4°. As the last residue of d1 (C115) participates in one of the three conserved disulfide bridges (C2C6) and the first residue of d2 (d116) is hydrogen bonded to Y176 of the same domain, there are no additional linker residues that are not assigned to either d1 or d2. The d2 helix α1 plays a crucial role in the formation of the d1_d2 interface by donating three residues to it (I173, Y176, F177), which are mostly interacting with other hydrophobic residues of the d1 C-terminus (F112, V114). Other N-terminal residues of the d2 domain (W117, H118) interact mostly with the C2C6 disulfide bridge of the d1 domain. Since due to the C-terminal tip (d1) / N-terminal tip (d2) arrangement of the d1_d2 interface the overall arrangement is asymmetric, the CC(1)-W(2)-PP(3)-R(4)-W(5)-QQ)6)-CC(7) layer structure in d2 is on the inward face of the overall d1_d2 V-arrangement, whereas the same layer structure in d2 is on the outwards face.

To validate our structural findings, we used Small Angle X-ray Scattering (SAXS). Investigation of the distance distribution *P*(*r*), maximum particle dimension and the generation of a dummy residue models confirmed the V-shape arrangement of the THSD7A d1_d2 structure **(Figure 6, Supplementary Figure S3, Supplementary Table S2)**. The additional distinct local distance maximum in the *P*(*r*) distribution near *r* = 4 nm, which is characteristic for dumbbell-shaped or modular/domain-separated molecules, most likely reflects a distance range of the distally folded β-sheet structures by domains d1 and d2 within the V-shaped arrangement **(Figure 6B)**. While the two tips of the V-shape distal ends reach a maximum distance of about 6 nm, the centers of scattering mass that are relevant for SAXS-based estimates are in approximate 4 nm distance, thus well supporting this interpretation. By generating a hybrid model, in which sequence segments missing in the d1_d2 crystal structure were added from the available AF2 model **(Supplementary Figure S2)**, the *χ* ^2^ value describing the fit with the experimental SAXS curve could be improved from 4.2 to 1.1. When using a d1_d2 (d116A, H118A, Y176A) variant termed d1_d2(AAA), in which key d1_d2 interface interactions were removed, the SAXS profile markedly changed **(Figures 2B and 6A)**. The radius of gyration (*R*_*g*_) increased from 2.58 nm to 2.83 nm, and similarly the maximum distance (*D*_*max*_) value increased from 9.5 nm to 11.5 nm. The resulting dummy residues model indicates opening up of the V-shaped arrangement of *wt* d1_d2 towards an extended arrangement, one would expect from a flexible linker **(Figure 6C)**. The disappearance of the second local intensity maximum in the distance distribution *P*(*r*) plot most likely reflects the loss of a defined d1_d2 arrangement **(Figure 6B)** whereby the mutant is best described as an ensemble of structural states in solution (**Supplementary Figure 2**).

**Figure 6:**
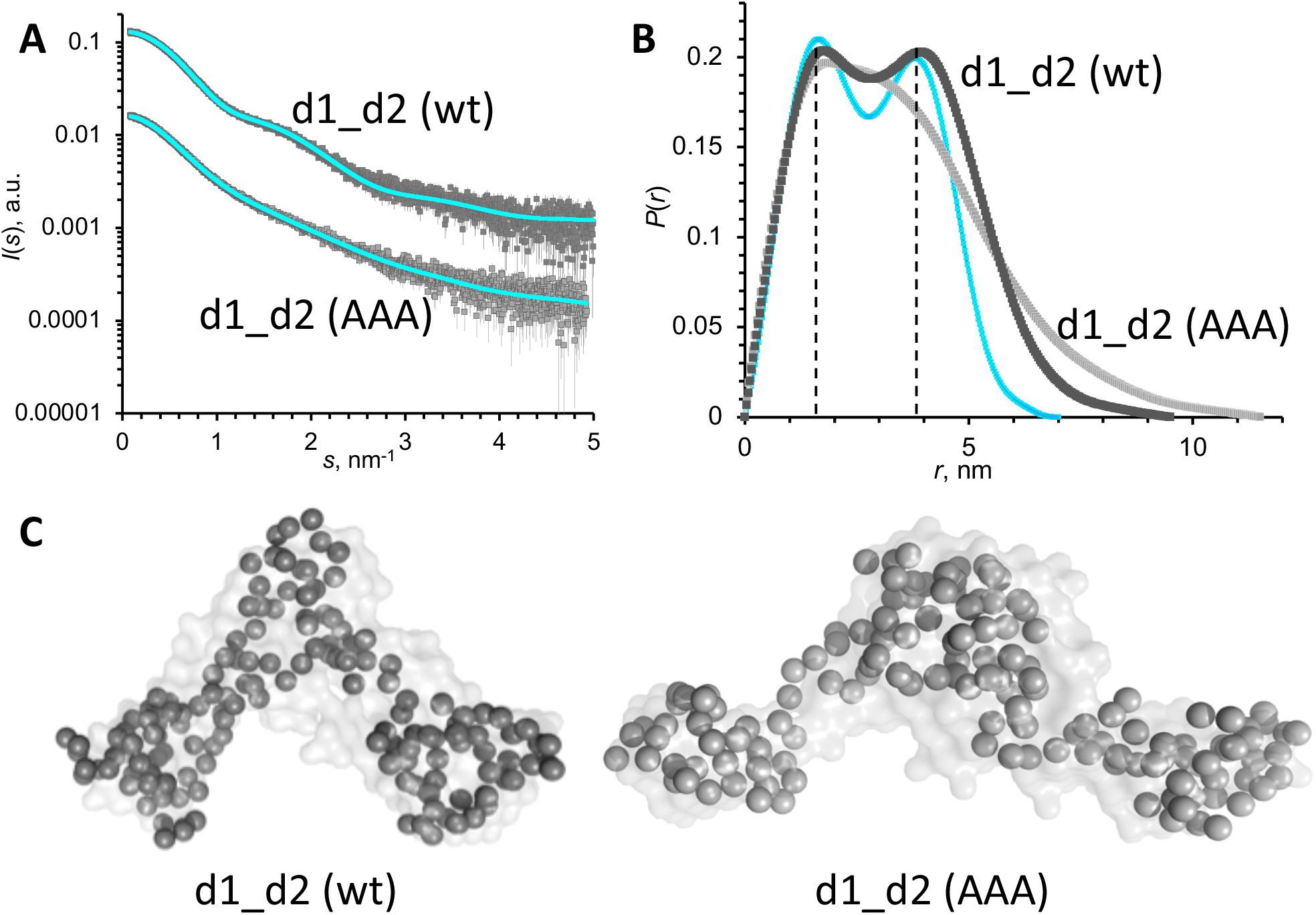
THSD7A d1_d2 structural validation by SAXS. A, SAXS curves of d1_d2(wt) and d1_d2(AAA) in dark grey and light grey, respectively. Model fits are in cyan. B, *P*(*r*) plot of d1_d2(wt) and d1_d2(AAA), indicating a second distance maximum at about 4 nm distance. The SAXS curve calculated from the d1_d2 crystal structure is also shown (cyan). These maxima are characteristic for dumbbell-type molecular shapes with defined, modular domain arrangements. C, dummy residue models of d1_d2(wt, left) and d1_d2(AAA, right). Color codes: d1_d2(wt), dark grey; d1_d2(AAA), light grey.

### Type 1A TSR fold of THSD7A as model for other THSD7A TSR domains

We used our structural findings of conserved structural and sequence features of domains THSD7A d1 and d2 to assess the likelihood to find this fold in other THSD7A domains that were predicted to belong to the type 1 TSR fold (Stoddard et al., 2019). To this end, we used the conserved CC(1)-W(2)-PP(3)-R(4)-W(5)-QQ)6)-CC(7) layer structure as reference for comparison **(Figure 3, Table 1)**. Interestingly, there is low conservation of this pattern in other relevant THSD7A domains. The layer 3 PP pattern is only found in D4, d10 and d16. Since in D4 the first disulfide bridge of layer 1 and the conserved tryptophan of layer 2 are missing, d10 and d16 remain the most likely candidates to be similarly folded as type 1A TSR domains. The only observed deviation is in the QQ layer 6, where replacement by other hydrophilic residues are found. As the side chain of this position points towards the solvent exterior changes in this residue position are probably less critical for fold definition. As in all other domains (d6, d8, d12, d14, d18, d20) the layer 3 PP motif is missing or only partly present it remains uncertain whether these domains are folded similarly to the type 1A fold found in d1 and d2.

**Table 1:**
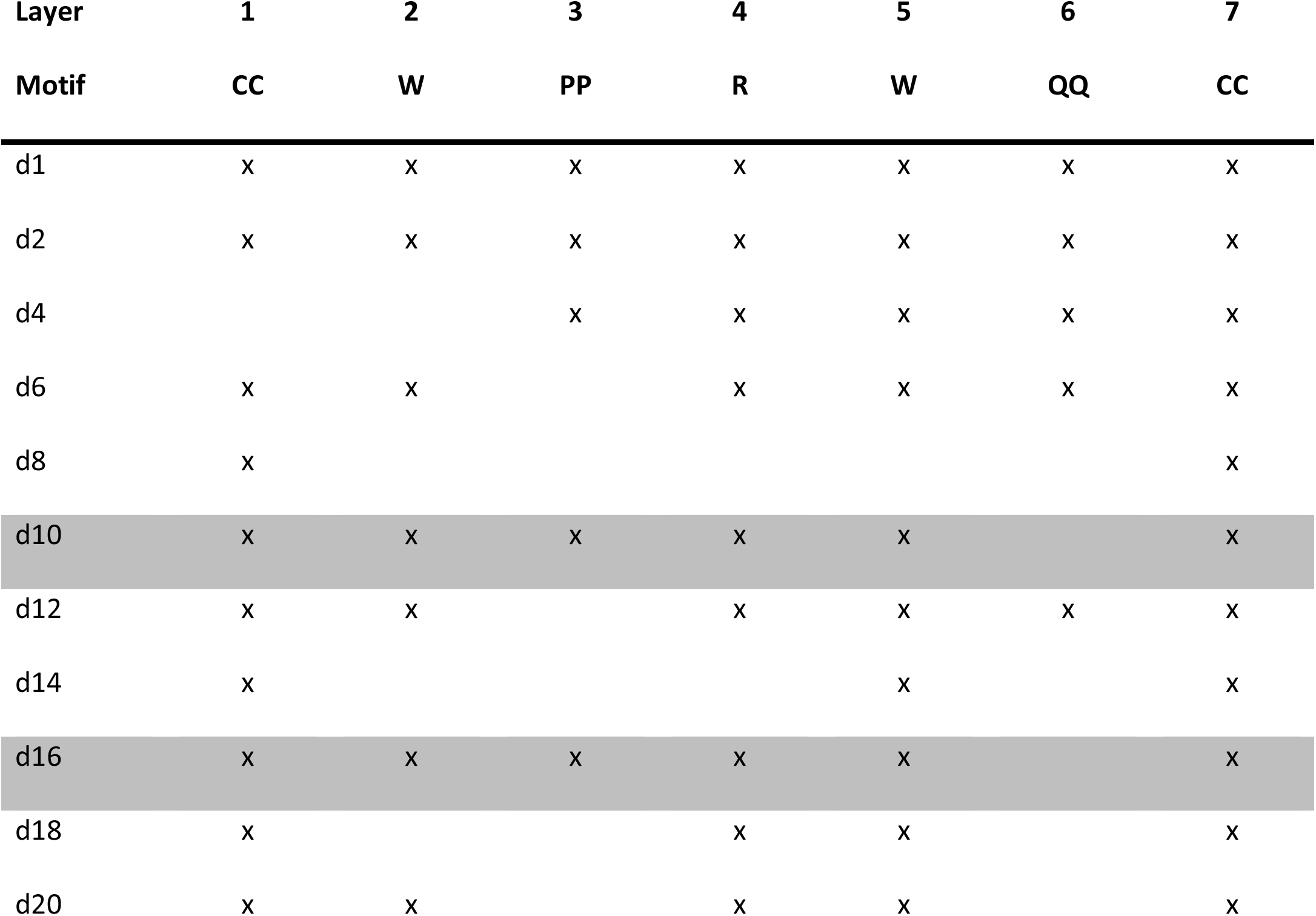
CC-W-PP-R-W-QQ-CC layers in predicted THSD7A TSR domains with type 1 TSR disulfide pattern. THSD7A domains that are most likely to adopt a type 1A TSR fold (d10, d16) are highlighted in grey.

An open question in how neighboring domains in THSD7A could be arranged. This question is relevant as kinked domain/domain arrangements would be required for a compact overall THSD7A structure. Since the majority of next neighbor THSD7A TSR domain connections are very short (Seifert et al., 2018, Stoddard et al., 2019), it is likely that common interfaces of neighboring domains are formed with well-defined domain/domain arrangements. As many of these short linkers contain a proline residue (Seifert et al., 2018), it will be of particular interest to investigate their role in potentially defining any specific TSR/TSR domain arrangements in THSD7A. However, since the V-shaped arrangement observed for d1_d2 is the only one between two type 1A TSR domains, it is unlikely that this could serve as blueprint for other domain/domain arrangements. The modified disulfide pattern of type 2 TSR domains, presenting the remaining half of TSR domains in THSD7A, links the very N-terminus of each domain by an additional disulfide bridge (C1C4), which does not exist in type 1 TSR domains. The presence of these bridges could play an additional crucial role in defining next neighbor TSR/TSR domain arrangements in THSD7A. It is also worth noting that despite the well-defined V-shaped arrangement of d1_d2, in contrast to most other connecting TSR/TSR linkers there is no proline in the d1/d2 linker.

## Materials and Methods

### Design and expression of mouse THSD7A variants

For our experiments we designed 11 fragments of mouse THSD7A (mTHSD7A) d1_d2 (Ala-37 to Gln-181), d2_d3 (Trp-106 to Glu-236), d3_d5 (Lys-182 to Ala-412), d6_d7 (Thr-413 to Tyr-563), d8_d9 (Asp-564 to Thr-684), d10_d11 (Val-685 to His-820), d12_d13 (Ser-821 to Asp-888), d14_d15 (Lys-889 to Asn-1084), d16_d17 (Gln-1085 to Tyr-1208), d18_d19 (His-1209 to Tyr-1329) and d20_d21 (Arg-1330 to Tyr-1463), reassembling the whole extracellular part of mTHSD7A.

Additionally, a construct comprising all mTHSD7A extracellular domains with an N-terminal strep-tag was designed (d1_d21). All variants were generated by PCR, with cDNA of a full-length, flag-tagged mTHSD7A construct serving as the PCR template (Origene).

For immunization d1_d2 and d1_d21 were cloned into the eukaryotic expression vector pCSE2.5 (provided by Thomas Schirrmann, Braunschweig, Germany), resulting in C-terminal hexa-histidine tagged constructs. For ELISA analysis all variants were cloned into the eukaryotic expression vector pDSG-IBA104 (IBA-Lifescience), resulting in N-terminal twin strep tagged proteins. Full sequencing validated the accuracy of all constructs.

For protein expression, HEK293F cells were cultured in 30 ml of Freestyle 293 serum-free medium (Gibco). For transfection of HEK293F cells, a polyethylenimine (PEI) based method (Polyscience) was used. For each approach, 126 μL (40 μg) of PEI was mixed with 124 μL water and 250 μL NaCl (300 mM) was added. 10 μg plasmid DNA in 250 μL water was mixed with the same amount of NaCl in a separate tube. Subsequently, solutions were slowly mixed, vortexed and incubated for 30 minutes. The solution was then added to the cells. After 24 h, the cells received 250 μL Freestyle 293 medium supplemented with 20% tryptone. Five days later, cells were collected using centrifugation at 300 g and the supernatant was centrifuged again at 14,000 g for 10 minutes. Hexa-histidine tagged mTHSD7A fragments were purified under native conditions using Ni-NTA resin (Thermo Scientific) applying the batch method according to the manufacturer’s instructions. After purification, the buffer was exchanged to PBS using ZEBA-spin columns (Thermo Scientific).

Twinstrep tagged mTHSD7A variants were purified under native conditions using strep-tactin resin, following the manufacturer’s instructions (Strep-Tactin XT, IBA-lifescience). All expressions and purifications were validated by Western blot and/or Coomassie staining.

### Animal experiments

Animal experiments were performed according to national and institutional animal care and ethical guidelines and were approved by the Veterinarian Agency of Hamburg and the local animal care committee (registration numbers 114/18 and 002/19). Wild-type male BALB/c were bred in the animal facility of the University Medical Center Hamburg-Eppendorf. Animals had free access to water and standard animal chow. Mice lacking THSD7A (*Thsd7a*^*−/ −*^ mice) were generated as described (Seifert et al., 2023).

To obtain mouse serum containing anti THSD7A antibodies *Thsd7a*^*−/ −*^ mice were immunized with the mTHSD7A fragments d1_d2 or d1_d21. Mice received a total of 3 immunizations: 20 μg of protein per fragment were mixed with an equal volume of complete Freund’s adjuvant and injected subcutaneously, followed by two boost immunizations of 20 μg per fragment in an equal volume of incomplete Freund’s adjuvant after 3 and 5 weeks. Control mice received the equal amounts of adjuvant diluted in PBS. 2 weeks after the last immunization the mice were euthanized for the collection of blood.

For the purification of IgG from mouse sera, we used Protein G columns (Thermo Scientific). Columns were washed with 25 ml PBS before serum application. After complete drainage, columns were washed with 20 ml PBS and IgG was eluted in 3 steps, using 4.5 ml 100 mM glycine (pH 3) each time. To neutralize the eluate was collected in Tris (pH 9). The buffer was exchanged for PBS by size exclusion chromatography on PD-10 columns (GE Healthcare). Purification of IgG was verified using SDS-PAGE and Coomassie blue staining. Elution fractions containing highest concentrations of IgG were pooled and concentrated using Amicon Ultra-15 centrifugal filters (Millipore). The final concentration of purified mouse IgG was determined with a Bradford assay. Two 15-week-old male BALB/c mice were injected intravenously with 1.5 mg of anti-d1_d2, anti-THSD7A (d1_d21) or control IgG, respectively.

Development of proteinuria was monitored using metabolic cages 1 day before injection of IgG and on day 2 and 5 after injection. Mice were euthanized after 7 days for blood, urine and organ collection. Albumin content in urine was quantified using a commercially available ELISA system (Bethyl) according to the manufacturer’s instructions. The urine albumin values were standardized against urinary creatinine values, as determined according to Jaffé (Hengler Analytik) of the same sample.

### Anti-THSD7A ELISA

Microplates (Sarstedt) were coated with 100 ng of different mTHSD7A variants per well in carbonate-bicarbonate buffer over night at 4 °C, washed 3 times with Tris buffered saline with Tween 20 (pH 8), and blocked with BSA in Tris buffered saline (pH 8) for 30 minutes. Mouse sera were diluted 1:100 in Tris buffered saline with BSA (pH 8), and incubated for 2 h followed by 3 washing steps. Next, samples were incubated with HRP-conjugated goat anti-mouse IgG antibody at 1:5000 ratio (Jackson Immuno Research Laboratories), diluted in Tris buffered saline with BSA (pH 8) for 1 h and subsequently washed for 4 times. Then, the TMB substrate (Aviva Systems Biology) was added for 2 minutes. The reaction was stopped with 1 M H_3_PO_4_. All incubation steps were carried out at room temperature (RT). The optical density was read at 450 nm using an automated spectrophotometer (Ultra-microplate reader, Bio-TEK instruments).

### Immunofluorescence and electron microscopy

Paraffin sections (3-4 μm) of formalin fixed mouse kidneys were deparaffinized and rehydrated. Antigen retrieval was obtained by boiling in Dako Target Retrieval (pH 9) (Dako) for 30 min in a steamer at 98°C, or by digestion with 5 μg/mL protease XXIV (Sigma-Aldrich) for 15 min at 37°C. Unspecific binding was blocked with blocking buffer (5% horse serum (Vector Laboratories) supplemented with 0.05% Triton X-100 (Sigma-Aldrich) in PBS) for 30 min at RT before incubation at 4°C overnight with primary antibodies in blocking buffer. Staining was visualized with affinity purified, fluorochrome-conjugated secondary antibodies at a ratio of 1:200 (Jackson ImmunoResearch Laboratories, Invitrogen) for 30 min at RT in blocking buffer. Nuclei were counterstained with DRAQ5 at ratio of 1:1000 (Cell Signaling), DAPI at ratio of 1:400 (Sigma-Aldrich) or Hoechst33342 at a ratio of 1:1000 (Sigma-Aldrich). Sections were mounted with Fluoromount-G (Invitrogen). Optical images were obtained using an inverted laser confocal microscope LSM800 (Zeiss).

For immunolocalization, the following antibodies were used: guinea pig nephrin at a ratio of 1:200 (Acris Antibodies BP5030), WGA-rhodamin at a ratio of 1:400 (Vector-Laboratories), goat THSD7A at a ratio of 1:200 (Santa Cruz Biotechnology) or rabbit THSD7A at a ratio of 1:200 (Sigma-Aldrich), C1q goat serum at a ratio of 1:400 (Complement Technology), C3 FITC goat anti-C3 at a ratio of 1:100 (Cappel), Cy2 donkey anti-mouse IgG H+L at a ratio of 1:200 (Jackson ImmunoResearch Laboratories).

Electron microscopic analyses were performed on kidneys that were fixed in 4% buffered paraformaldehyde. Tissue was post-fixed with 1% osmium in 0.1 M sodium-cacodylate buffer, stained with 1% uranyl acetate and embedded in epoxy-resin (Serva). Ultra-thin sections were cut (Ultramicrotome, Reichert-Jung) and contrasted with uranyl acetate in methanol followed by lead citrate. Micrographs were generated with a JEM 1010 transmission-electron microscope (JEOL).

### Expression and purification of d1_d2 for structural analyses

The gene encoding human THSD7A domains d1_d2 (UNIPROT Q9UPZ6, residues A48–Q192) was sub-cloned into the eukaryotic expression vector pDSG-IBA104 (IBA Lifesciences, Göttingen, Germany). It positioned a BM40 secretion signal, Twin-Strep-tag and Human rhinovirus (HRV) 3C protease cleavage site (WSHPQFEKGGGSGGGSGGSAWSHPQFEK) at the N-terminus of d1.

For expression of recombinant d1_d2 protein, HEK293F cells (Thermo Scientific) were transiently transfected at a cell density of 20 × 10^6^ cells/mL. Transfection was performed with d1_d2 pDSGIBA104 plasmid DNA and PEI (Polysciences) in a 1:3 (w/w) ratio, respectively. After adding the DNA/PEI mixture, the cell culture was cultivated for 1 h. Next, the fresh medium (FreeStyle 293 expression medium (Thermo Scientific) was added to dilute the cells to a density of 1 × 10^6^ cells/mL. After 24 h post-transfection, the cell culture was supplemented with valproic acid to a final concentration of 3.75 mM. The transfected cells were grown in suspension while shaking in a humidified incubator at 37 °C and 8% CO_2_. The cell medium containing the secreted protein was harvested after 5 days by centrifugation at 100 g for 5 minutes at RT. 1.6 mL/L BioLock (IBA Lifesciences, Göttingen, Germany) was added to the supernatant and the pH was adjusted to 8.0 with 50 mM Tris pH 9.0. After incubation on ice for 30 min, the supernatant was clarified by centrifugation at 10,000 g. Subsequently, the tagged d1_d2 was purified from the supernatant by affinity chromatography using Strep-Tactin resin (IBA Lifesciences, Göttingen, Germany).

Following overnight incubation at 4 °C with Strep-Tactin resin, the resin was washed with a buffer comprising 50 mM Tris (pH 8.0), 150 mM NaCl. Tagged d1_d2 was eluted from resin with a buffer comprising 50 mM Tris (pH 8.0), 150 mM NaCl, and 5 mM desthiobiotin). The N-terminal strep tag was removed by in-house purified 3C protease in a 1:30 (protein:3C) ratio, followed by overnight incubation at 4 °C. Finally, the d1_d2 was further purified by size exclusion chromatography (SEC) using a column Superdex® 75 Increase 10/300 GL (Cytiva), pre-equilibrated with a buffer comprising 20 mM HEPES (pH 7.4), and 100 mM NaCl. Protein purity was evidenced by a sharp, mono-disperse elution profile and as a clear band on SDS-PAGE.

### X-ray crystallography

A sparse, sitting drop vapor diffusion crystallization trials was set up using the Hampton Index screen (Hampton research) with purified d1_d2 at a concentration of 5.3 mg/mL in CrystalDirect 96-well plates. Equal volumes 1:1 of protein and precipitant were combined. Crystals for structural analysis were grown at 4 °C from 0.1 M sodium chloride, 0.1 M bis-tris (pH 6.5), 1.5 M ammonium sulphate. Suitable crystal was mounted and washed with the reservoir solution several times and cryoprotected through brief soaking in paratone-N oil before being flash frozen in liquid nitrogen. X-ray diffraction data were collected at beamline P14 at the PETRA III synchrotron ring (EMBL/DESY, Hamburg, Germany) using an EIGER 16M detector. Diffraction images were processed using XDS (Kabsch, 2010). The d1_d2 structure was solved by molecular replacement using Phaser (McCoy et al., 2007) and models of d1 and d2 generated by AlphaFold2 (Q9UPZ6) as search models. Residues A48–Y59, G91–T93 of d1 and V135-E150 of d2 were deleted from the search models. The resulting structure was subjected to several rounds of refinement and model building using phenix refine (Adams et al., 2010), REFMAC5 (Murshudov et al., 2011) and Coot (Emsley et al., 2010). There was no electron density for amino acids A48–A53 at the N-terminus of d1, S137–E145 in the AB loop of d2 or Q192 at the C-terminus of d2. The final model had R_work_ and R_free_ of 22.9 % and 28.1 % respectively. Figures were generated in PyMOL (Schroedinger). Coordinates and structure factors have been deposited in the PDB with the accession code 8OXR. Angles between the domains d1 and d2 structures were measured using best-fit lines to the domain coordinates determined using Principal Component Analysis.

### Size exclusion chromatography small-angle X-ray scattering (SEC-SAXS)

SAXS data of wild-type THSD7A d1_d2 (d1_d2, wt) and the triple alanine mutant (d1_d2, AAA) were measured at beamline P12 at the PETRAIII synchrotron ring (EMBL/DESY, Hamburg, Germany), coupled with size exclusion chromatography (Blanchet et al., 2015). Samples were injected onto a Superdex® 75 Increase 5/150 column (Cytiva) pre-equilibrated with 25 mM Tris, 150 mM NaCl, 5 mM NaNO_3_ (pH 7.5) at a flow rate of 0.35 mL/min at 20 °C, where the continuously flowing eluate was passed directly to the beam line. SAXS intensities were measured from the column eluate as a set of sequential 0.24–0.25 s images using a PILATUS 6M detector (Hajizadeh et al., 2018). Subsequent azimuthal averaging of the intensities to generate sets of reduced 1D-data frames was performed using standard beam line protocols(Franke et al., 2012). Additional SEC-SAXS data processing was performed using *CHROMIXS* (Panjkovich and Svergun, 2018) and modules of the *ATSAS 3*.*0* software package (Manalastas-Cantos et al., 2021) to generate the background subtracted and averaged SAXS profiles of d1_d2 variants in solution.

Shapes of the d1_d2 variant were reconstructed using the dummy-residue approach implemented in *GASBOR* (Svergun et al., 2001). Five individual *GASBOR* models that fit the respective SAXS profiles underwent spatial alignment with *DAMSEL* and *DAMSUP*, to generate an overall/global representation of the average structural disposition of either protein in solution (Volkov and Svergun, 2003). Further atomistic representations were built using a combined approach, taking the d1_d2 crystal structure (8OXR) and adding the atomic coordinates of amino acids from missing regions, using an AlphaFold-predicted model template (Q9UPZ6) (Varadi et al., 2022). The resulting hybrid atomic model was built using a structure-structure alignment procedure in PyMOL (Schroedinger), by combining atomic coordinates of the crystal structure with the AlphaFold atomic coordinates of missing regions. Additional modelling of the d1_d2(AAA) variant was performed using Ensemble Optimization Method (Bernadó et al., 2007, Tria et al., 2015) by taking the initial d1_d2(wt) hybrid model and generating a pool of 10,000 randomly oriented structures, and selecting volume-fraction weighted sub-ensembles from the initial pool based on the fit to the SAXS data. The model/data-fit evaluations for both d1_d2(wt) and d1_d2(AAA) ensembles were evaluated using the reduced *χ*^2^ test and Correlation Map (CorMap) *P*-value (Franke et al., 2015). All data and models have been deposited to the Small Angle Scattering Biological Data Bank (Kikhney et al., 2020) under the accession codes:d1d2(wt), SASDRZ6; d1_d2(AAA), SASDR27.

## Abbreviations

AF2: AlphaFold2
MN: Membranous Nephropathy
r.m.s.d.: root-mean-squares-deviation
THSD7A: Thrombospondin type-1 domain-containing 7A
TSR: Thrombospondin repeat

## Acknowledgements

SAXS and X-ray diffraction data were collected at beamlines P12 and P14 operated by the EMBL Hamburg Unit at the PETRA III storage ring (DESY, Hamburg, Germany), respectively. We acknowledge technical support by the Sample Preparation Characterization facility at EMBL Hamburg. We thank Sihyun Sung (EMBL), Barbara Ramsak (EMBL), Mira Schiffhauer (EMBL) and Sofia Mortensen (DESY, Hamburg, Germany) for technical and scientific support. Gunter Zahner (UKE) is thanked technical assistance.

## References

Adams, P. D., Afonine, P. V., BunkóCzi, G., Chen, V. B., Davis, I. W., Echols, N., Headd, J. J., Hung, L. W., Kapral, G. J., Grosse-Kunstleve, R. W., Mccoy, A. J., Moriarty, N. W., Oeffner, R., Read, R. J., Richardson, D. C., Richardson, J. S., Terwilliger, T. C. & Zwart, P. H. 2010. PHENIX: a comprehensive Python-based system for macromolecular structure solution. Acta Crystallogr D Biol Crystallogr, 66, 213–21.

Beck, L. H., JR., Bonegio, R. G., Lambeau, G., Beck, D. M., Powell, D. W., Cummins, T. D., Klein, J. B. & Salant, D. J. 2009. M-type phospholipase A2 receptor as target antigen in idiopathic membranous nephropathy. N Engl J Med, 361, 11–21.

Bernadó, P., Mylonas, E., Petoukhov, M. V., Blackledge, M. & Svergun, D. I. 2007. Structural characterization of flexible proteins using small-angle X-ray scattering. J Am Chem Soc, 129, 5656–64.

Blanchet, C. E., Spilotros, A., Schwemmer, F., Graewert, M. A., Kikhney, A., Jeffries, C. M., Franke, D., Mark, D., Zengerle, R., Cipriani, F., Fiedler, S., Roessle, M. & Svergun, D. I. 2015. Versatile sample environments and automation for biological solution X-ray scattering experiments at the P12 beamline (PETRA III, DESY). J Appl Crystallogr, 48, 431–443.

Crine, S. L. & Acharya, K. R. 2022. Molecular basis of C-mannosylation - a structural perspective. Febs j, 289, 7670–7687.

Emsley, P., Lohkamp, B., Scott, W. G. & Cowtan, K. 2010. Features and development of Coot. Acta Crystallogr D Biol Crystallogr, 66, 486–501.

Franke, D., Jeffries, C. M. & Svergun, D. I. 2015. Correlation Map, a goodness-of-fit test for one-dimensional X-ray scattering spectra. Nat Methods, 12, 419–22.

Franke, D., Kikhney, A. G. & Svergun, D. I. 2012. Automated acquisition and analysis of small angle X-ray scattering data. Nuclear Instruments and Methods in Physics Research Section A: Accelerators, Spectrometers, Detectors and Associated Equipment, 689, 52–59.

öDel, M., Grahammer, F. & Huber, T. B. 2015. Thrombospondin type-1 domain-containing 7A in idiopathic membranous nephropathy. N Engl J Med, 372, 1073.

Hajizadeh, N. R., Franke, D., Jeffries, C. M. & Svergun, D. I. 2018. Consensus Bayesian assessment of protein molecular mass from solution X-ray scattering data. Sci Rep, 8, 7204.

Herwig, J., Skuza, S., Sachs, W., Sachs, M., Failla, A. V., Rune, G., Meyer, T. N., Fester, L. & Meyer-Schwesinger, C. 2019. Thrombospondin Type 1 Domain-Containing 7A Localizes to the Slit Diaphragm and Stabilizes Membrane Dynamics of Fully Differentiated Podocytes. J Am Soc Nephrol, 30, 824–839.

Kabsch, W. 2010. XDS. Acta Crystallogr D Biol Crystallogr, 66, 125–32.

Kikhney, A. G., Borges, C. R., Molodenskiy, D. S., Jeffries, C. M. & Svergun, D. I. 2020. SASBDB: Towards an automatically curated and validated repository for biological scattering data. Protein Sci, 29, 66–75.

Kuo, M. W., Wang, C. H., Wu, H. C., Chang, S. J. & Chuang, Y. J. 2011. Soluble THSD7A is an N-glycoprotein that promotes endothelial cell migration and tube formation in angiogenesis. PLoS One, 6, e29000.

Manalastas-Cantos, K., Konarev, P. V., Hajizadeh, N. R., Kikhney, A. G., Petoukhov, M. V., Molodenskiy, D. S., Panjkovich, A., Mertens, H. D. T., Gruzinov, A., Borges, C., Jeffries, C. M., Svergun, D. I. & Franke, D. 2021. ATSAS 3.0: expanded functionality and new tools for small-angle scattering data analysis. J Appl Crystallogr, 54, 343–355.

Mccoy, A. J., Grosse-Kunstleve, R. W., Adams, P. D., Winn, M. D., Storoni, L. C. & Read, R. J. 2007. Phaser crystallographic software. J Appl Crystallogr, 40, 658–674.

Murshudov, G. N., SkubáK, P., Lebedev, A. A., Pannu, N. S., Steiner, R. A., Nicholls, R. A., Winn, M. D., Long, F. & Vagin, A. A. 2011. REFMAC5 for the refinement of macromolecular crystal structures. Acta Crystallogr D Biol Crystallogr, 67, 355–67.

Olsen, J. G. & Kragelund, B. B. 2014. Who climbs the tryptophan ladder? On the structure and function of the WSXWS motif in cytokine receptors and thrombospondin repeats. Cytokine Growth Factor Rev, 25, 337–41.

Panjkovich, A. & Svergun, D. I. 2018. CHROMIXS: automatic and interactive analysis of chromatography-coupled small-angle X-ray scattering data. Bioinformatics, 34, 1944–1946.

Polanco, N., GutiéRrez, E., Rivera, F., Castellanos, I., Baltar, J., Lorenzo, D. & Praga, M. 2012. Spontaneous remission of nephrotic syndrome in membranous nephropathy with chronic renal impairment. Nephrol Dial Transplant, 27, 231–4.

Resovi, A., Pinessi, D., Chiorino, G. & Taraboletti, G. 2014. Current understanding of the thrombospondin-1 interactome. Matrix Biol, 37, 83–91.

Schieppati, A., Mosconi, L., Perna, A., Mecca, G., Bertani, T., Garattini, S. & Remuzzi, G. 1993. Prognosis of untreated patients with idiopathic membranous nephropathy. N Engl J Med, 329, 85–9.

Seifert, L., Hoxha, E., Eichhoff, A. M., Zahner, G., Dehde, S., Reinhard, L., Koch-Nolte, F., Stahl, R. A. K. & Tomas, N. M. 2018. The Most N-Terminal Region of THSD7A Is the Predominant Target for Autoimmunity in THSD7A-Associated Membranous Nephropathy. J Am Soc Nephrol, 29, 1536–1548.

Seifert, L., Zahner, G., Meyer-Schwesinger, C., Hickstein, N., Dehde, S., Wulf, S., KöLlner, S. M. S., Lucas, R., Kylies, D., Froembling, S., Zielinski, S., Kretz, O., Borodovsky, A., Biniaminov, S., Wang, Y., Cheng, H., Koch-Nolte, F., Zipfel, P. F., Hopfer, H., Puelles, V. G., Panzer, U., Huber, T. B., Wiech, T. & Tomas, N. M. 2023. The classical pathway triggers pathogenic complement activation in membranous nephropathy. Nat Commun, 14, 473.

Stoddard, S. V., Welsh, C. L., Palopoli, M. M., Stoddard, S. D., Aramandla, M. P., Patel, R. M., Ma, H. & Beck, L. H., JR. 2019. Structure and function insights garnered from in silico modeling of the thrombospondin type-1 domain-containing 7A antigen. Proteins, 87, 136–145.

Svergun, D. I., Petoukhov, M. V. & Koch, M. H. 2001. Determination of domain structure of proteins from X-ray solution scattering. Biophys J, 80, 2946–53.

Tan, K., Duquette, M., Liu, J. H., Dong, Y., Zhang, R., Joachimiak, A., Lawler, J. & Wang, J. H. 2002. Crystal structure of the TSP-1 type 1 repeats: a novel layered fold and its biological implication. J Cell Biol, 159, 373–82.

Tomas, N. M., Beck, L. H., JR., Meyer-Schwesinger, C., Seitz-Polski, B., Ma, H., Zahner, G., Dolla, G., Hoxha, E., Helmchen, U., Dabert-Gay, A. S., Debayle, D., Merchant, M., Klein, J., Salant, D. J., Stahl, R. A. K. & Lambeau, G. 2014. Thrombospondin type-1 domain-containing 7A in idiopathic membranous nephropathy. N Engl J Med, 371, 2277–2287.

Tomas, N. M., Huber, T. B. & Hoxha, E. 2021. Perspectives in membranous nephropathy. Cell Tissue Res, 385, 405–422.

Tria, G., Mertens, H. D., Kachala, M. & Svergun, D. I. 2015. Advanced ensemble modelling of flexible macromolecules using X-ray solution scattering. IUCrJ, 2, 207–17.

Tucker, R. P. 2004. The thrombospondin type 1 repeat superfamily. Int J Biochem Cell Biol, 36, 969–74.

Varadi, M., Anyango, S., Deshpande, M., Nair, S., Natassia, C., Yordanova, G., Yuan, D., Stroe, O., Wood, G., Laydon, A., Zidek, A., Green, T., Tunyasuvunakool, K., Petersen, S., Jumper, J., Clancy, E., Green, R., Vora, A., Lutfi, M., Figurnov, M., Cowie, A., Hobbs, N., Kohli, P., Kleywegt, G., Birney, E., Hassabis, D. & Velankar, S. 2022. AlphaFold Protein Structure Database: massively expanding the structural coverage of protein-sequence space with high-accuracy models. Nucleic Acids Res, 50, D439–D444.

Volkov, V. V. & Svergun, D. I. 2003. Uniqueness of ab initio shape determination in small-angle scattering. Journal of Applied Crystallography, 36, 860–864.

Wang, C. H., Su, P. T., Du, X. Y., Kuo, M. W., Lin, C. Y., Yang, C. C., Chan, H. S., Chang, S. J., Kuo, C., Seo, K., Leung, L. L. & Chuang, Y. J. 2010. Thrombospondin type I domain containing 7A (THSD7A) mediates endothelial cell migration and tube formation. J Cell Physiol, 222, 685–94.

