## Supplementary Information for "Structure of the N-terminal didomain d1_d2 of the Thrombospondin type-1 domain-containing 7A" (from another version of this preprint: version 1, 2023-05-03)

**Supplementary Table 1: X-ray data collection and structure refinement statistics.**

| | |
|---|---|
| **X-ray data collection** | |
| Space group | C 2 2 $2_1$ |
| Unit cell dimensions a, b, c (Å) | 67.84, 108.71, 53.54 |
| X-ray wavelength (Å) | 0.976 |
| Resolution range (Å) | 57.56-2.30 (2.38-2.30) |
| Total reflections | 120354 (12055) |
| Unique reflections | 9135 (855) |
| Multiplicity | 13.2 (13.6) |
| Completeness (%) | 99.8 (100) |
| $R_{meas}$ | 0.19 (3.63) |
| Mean I/⍰(I) | 10.8 (1.0) |
| Wilson B factor ($Å^2$) | 48.4 |
| CC ½ | 1.0 (0.62) |
| **Structural refinement** | |
| Reflections used in refinement | 9111 |
| $R_{work}$ (%) | 22.6 |
| $R_{free}$ (%) | 27.9 |
| R.m.s.d. bond lengths (Å) | 0.0174 |
| R.m.s.d. bond angles ($^o$) | 2.187 |
| Ramachandran plot (%) | |
| Favored | 98.4 |
| Allowed | 1.6 |
| Outliers | 0.0 |
| Average B factor ($Å^2$) | 52.0 |
| Protein | 51.8 |
| Solvent | 56.4 |

### Supplementary Table 2: SAXS data statistics

| | | |
|---|---|---|
| **Sample details** | | |
| Organism | *Homo sapiens* | |
| *Scattering particle composition* | | |
| Protein | UniProt ID: Q9UPZ6, residues 48–192 | |
| *Sample environment/configuration* | | |
| Solvent composition | 25 mM Tris, 150 mM NaCl, 5 mM $NaNO_3$ (pH 7.5) | |
| Sample temperature | 20 °C | |
| In beam sample cell | 1 mm quartz capillary, in vacuum, continuous flow | |
| *Size Exclusion Chromatography SEC-SAS* | | |
| SEC column type | Superdex 75 Increase 5/150 | |
| SEC flowrate, mL/min | 0.35 | |
| | **D1D2 (wt)** | **D1D2(AAA mutant)** |
| Sample injection concentration, mg/ml* | 4.7 | 4.1 |
| Sample injection volume, µL | 37 | 50 |
| **SAXS data collection** | | |
| Data acquisition/reduction software | *SASFLOW, BECQUEREL, CHROMIXS,* ChemStation (Agilent) | |
| Source/instrument description or reference | P12 (EMBL/DESY, Hamburg, Germany), Pilatus 6M detector (Dectris) coupled to an 1260 Infinity II Bio-Inert HPLC system (Agilent). | |
| X-ray wavelength (energy) | 0.124 nm (10 keV) | |
| Measured $q$-range ($q_{min} – q_{max}$ nm$^{-1}$) | 0.0196–7.34 | 0.022–7.31 |
| Working $q$-range ($q_{min} – q_{max}$; Å$^{-1}$, nm$^{-1}$) | 0.081–5.0 | 0.084–5.2 |
| Intensity scale, normalized to beam transmission | arbitrary | absolute, cm$^{-1}$ (relative to water at 20 °C) |
| Exposure time(number of exposures) | 0.25 s (48) | 0.24 s (91) |
| **SAXS-derived structural parameters** | | |
| Methods/Software | *ATSAS 3* | |
| $I(0) \pm \sigma$ | 2672.3 (3.5) a.u. | 0.01614 (0.00003) cm$^{-1}$ |
| $R_g \pm \sigma$ (nm) | 2.54(0.01) | 2.68(0.01) |
| *min* < $qR_g$ < *max* limit (data point range) | 0.21–1.29 (23–176) | 0.24–1.28 (25–164) |
| Linear fit correlation, $R^2$ | 0.996 | 0.993 |
| *PDDF/P(r) analysis* | | |
| $I(0) \pm \sigma$ | 2679(3.9) arbitrary | 0.01632(0.00003) cm$^{-1}$ |
| $R_g \pm \sigma$ (nm) | 2.58 (0.01) | 2.83 (0.01) |
| $D_{max}$ (nm) | 9.5 | 11.5 |
| $q$-range (nm$^{-1}$) | 0.081–5.0 | 0.089–5.0 |
| $P(r)$ fit, $\chi^2$/CorMap-$P$ $^+$ | 0.996 / 0.351 | 1.05 / 0.836 |
| **Particle size estimates** | | |
| Methods/Software | *ATSAS 3* | |
| Porod volume, $V_p$, nm$^3$) | 28 | 19 |
| *Molecular weight (M) estimates, kDa* | | |
| From amino acid sequence*, monomer | 16.7 | 16.4 |
| From Bayesian inference (0.9 credibility interval) $^{++}$ | 16 (15–17) | 14 (13–15) |
| **Modelling** | | |
| *Shape modelling methods* | *GASBOR* | |
| $q$-range for fit ($q_{min} – q_{max}$; nm$^{-1}$) | 0.081–5.0 | 0.089–5.0 |
| Symmetry/anisotropy assumptions | P1, prolate | P1, prolate |
| Number of model reconstructions | 5 | 5 |
| Model fit, $\chi^2$/CorMap-$P$ $^+$ | 1.06 / 0.013 | 1.07 / 0.836 |
| *Atomistic modelling methods* | X-ray crystal structure (PDB: 8OXR) /AlphaFold hybrid template | |

| | | |
|---|---|---|
| Software | *CRYSOL* | *EOM* |
| $q$-range for fit ($q_{min} - q_{max}$; $nm^{-1}$) | 0.081–5.0 | 0.084–4.9 |
| Symmetry/anisotropy assumptions | P1 | P1 |
| Model fit, $\chi^2$/CorMap-$P$ $^+$ | 1.09 / 0.193 | 1.02 / 0.818 |
| **Data and model deposition** | | |
| SASBDB IDs | SASDRZ6 | SASDR27 |

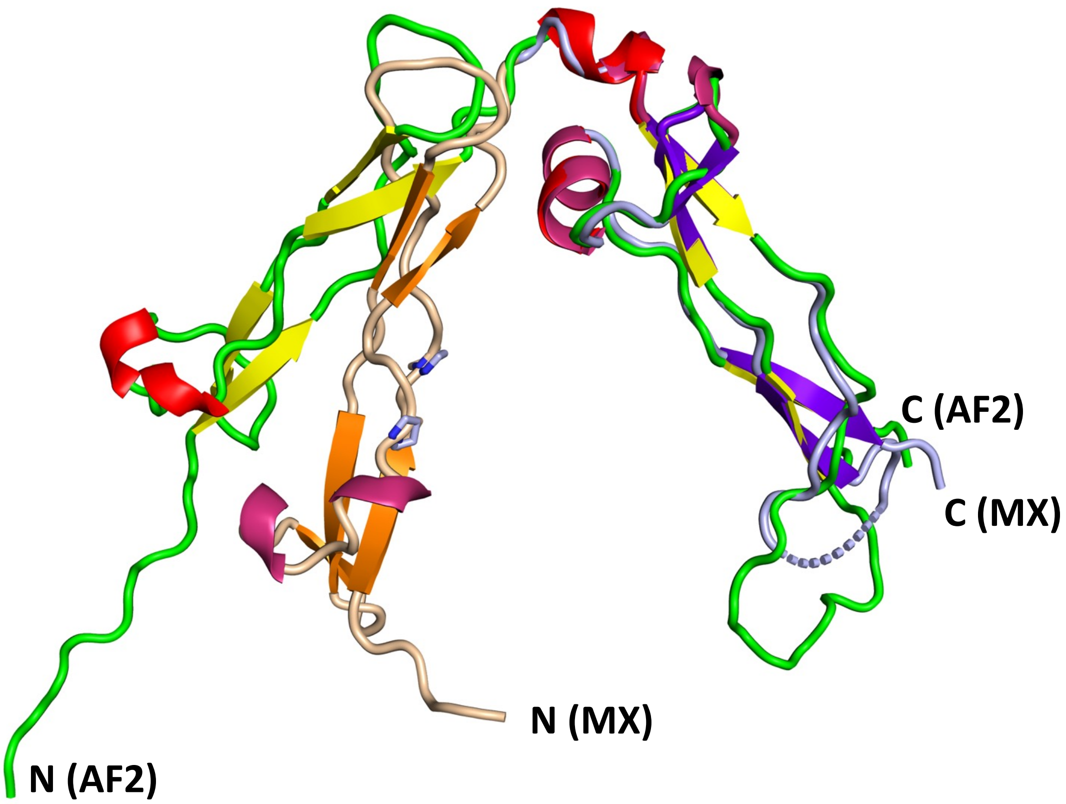


**Superposition of the AF2 model of THSD7A domains D1 and D2 (Q9UPZ6) onto the experimental crystal structure of the same construct (MX), using the D2 domain as basis for superposition.** Color codes of the MX structure are as in Figure 1. Color codes AF2 model: predicted β-strands, yellow; predicted α-helices, red; loop segments, green.

**Supplementary Figure 1**

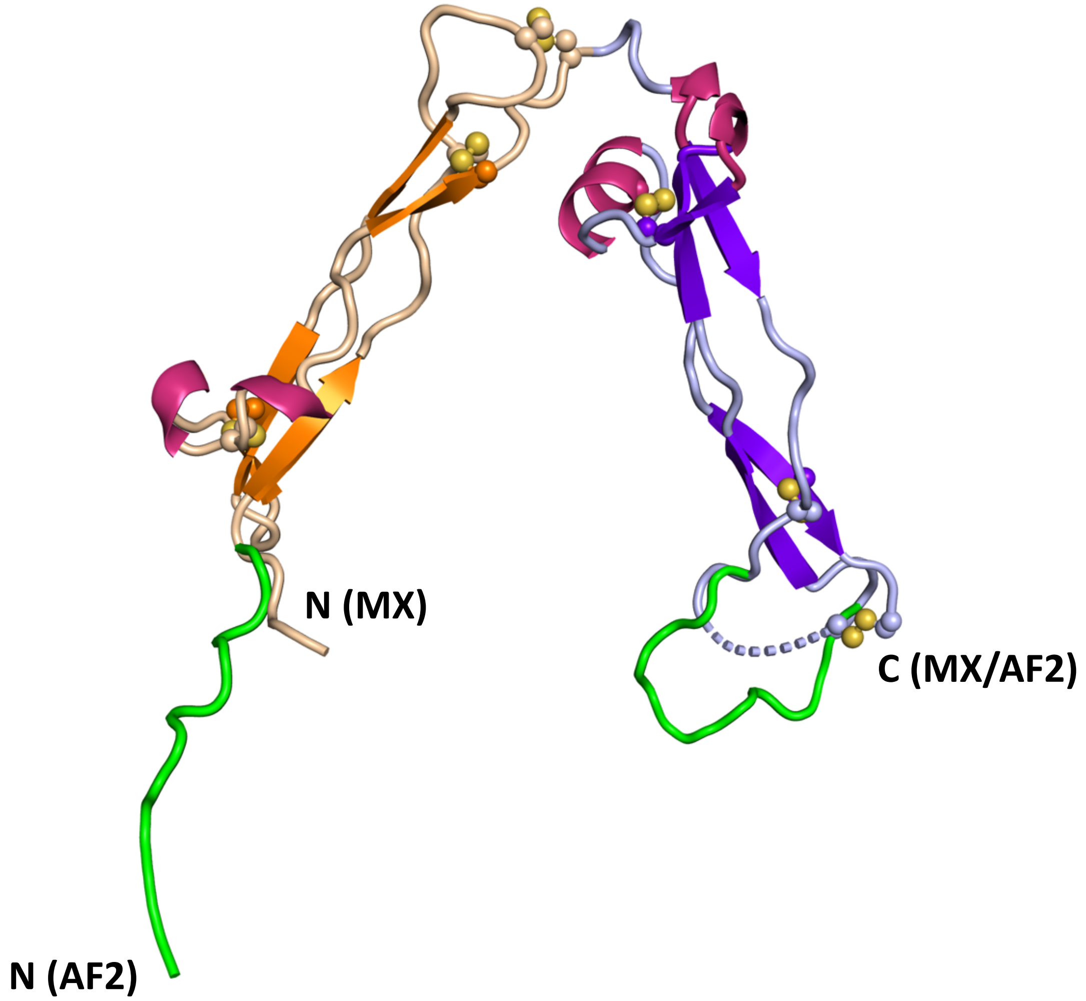


**Crystal structure / AF2 hybrid model of THSD7A D1D2.** AF2 model extensions include residues 46-58 (D1 N-terminus) and residues 134-147 (disordered AB loop of D2). Colors of the THSD7A D1D2 crystal structure are as in Figure 1. AF2 extensions are in green.

**Supplementary Figure 2**

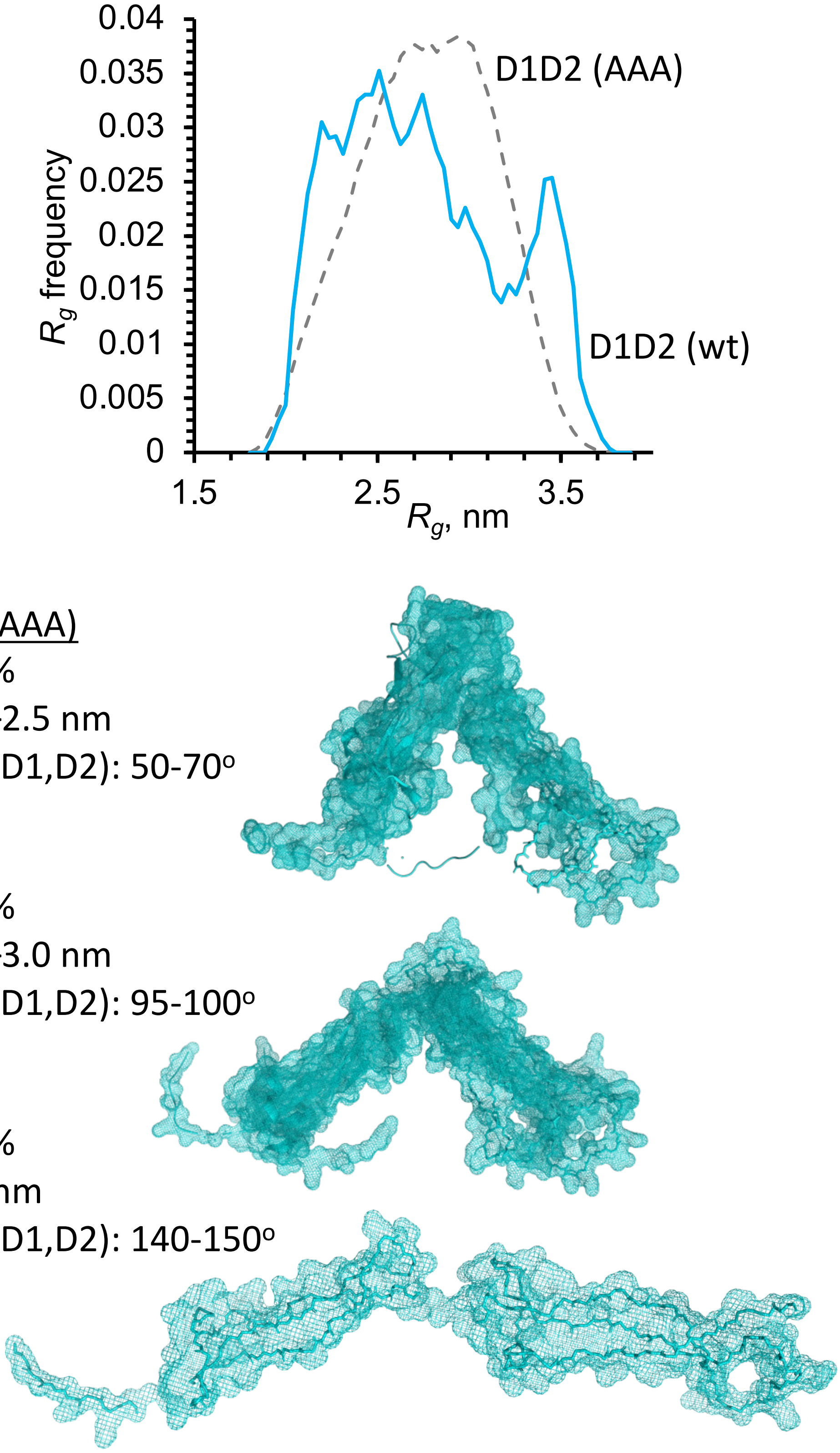


**Volume-fraction weighted ensemble of structural states of D1D2(AAA).** A, *Rg* frequency distribution; B, resulting ensemble of D1D2(AAA) models.

**Supplementary Figure 3**
